# Toxic effects of elevated bile acid levels on fetal myocardium in intrahepatic cholestasis of pregnancy, a retrospective study from a neonatal perspective

**DOI:** 10.1101/2022.05.16.492192

**Authors:** Junnai Wang, Wei Shi, Weiwei Lun

**Affiliations:** The Third Affiliated Hospital of Zhengzhou University, Zhengzhou, He’ nan 450052, China

**Keywords:** intrahepatic cholestasis of pregnancy, cardiac dysfunction

## Abstract

**Introduction:** Intrahepatic cholestasis of pregnancy (ICP) is a liver disease which may lead to a sudden fetal death. Previous studies have suggested that the fetal accident may be related to their cardiac dysfunction. However, the relationship between fetal cardiac dysfunction and their maternal bile acid levels is not clear. The objective was to clarify the relationship from a neonatal perspective and to furtherly make clear the after-effect by analysing the cardiac parameters of the older neonates.

**Materials and methods:** In this case-control study, patients and their neonates, managed between 10 September 2018 and 30 June 2021 at a Chinese university hospital center, were divided into severe ICP group, mild ICP group and control gestational group. The maternal bile acid levels and the cardiac paramerers of one-day-old neonates and five-day-old neonates were analysed, respectively.

**Results:** The specific-myocardial enzyme(CK-MB) and left ventricular fraction shortening(FS) of neonates showed significant differences between ICP group and control group, and were meaningfully correlated with maternal bile acid levels. However, There was no significant difference in cardiac injury parameters of older neonates between the ICP and control groups.

**Conclusions:** The elevated maternal bile acid levels can lead to fetal myocardial injury and the injury can be recovered after removel from high concentrations of bile acid.

## Introduction

Intrahepatic cholestasis of pregnancy (ICP) is the most common gestational liver disease and a pregnancy-specific disease. Clinically, it is characterized by itchy skin and elevated serum bile acid in pregnant women, and can lead to preterm birth, amniotic fluid fecal contamination, fetal distress, and unpredictable fetal death^[1]^. At present, there is no unified diagnosis and treatment for this disease, but the consensus view is that total bile acid levels are closely related to perinatal outcomes. The incidence of intrauterine fetal death in pregnant women with severe ICP can reach 9.5%–15.4%, which is 3 times higher than that in normal pregnancy. However, the mechanism of fetal death has not been clarified, and it is believed that cardiac factors may be an important cause of fetal death^[2]^.

In recent years, many studies have targeted fetal heart injury with ICP^[3]^. At autopsy, fetal stillbirths with ICP have exhibited spot-like hemorrhages in the pericardium, unclear structure of longitudinally striated myocardium, vacuolar degeneration, and spot-like necrosis^[4]^. Based on these pathological manifestations, studies on fetal myocardial injury mainly investigate the myocardial enzyme spectrum, cardiac function, and electrocardiogram (ECG)^[5-7]^. For example, umbilical cord blood from fetuses of normal pregnant women or pregnant women with ICP have been used to study the relationship between serum total bile acid levels and myocardial enzymes or troponin-I levels^[8]^. It was found that the levels of lactate dehydrogenase (LDH), α-hydroxybutyrate dehydrogenas(HBDB), and creatine kinase (CK) were significantly higher in fetuses of the severe ICP group than those in the mild ICP group, and troponin-I was higher in the severe ICP group than that in normal group^[9]^. Examination of fetal ECG in normal and ICP groups showed that the PR interval was prolonged in ICP patients, suggesting that cardiac conduction function had changed. Also, the fetal PR interval was longer in severe compared with mild ICP. Some studies found that the Tei index, isovolumic systolic time, and isovolumic diastolic time were significantly higher in fetuses of the ICP group versus the control group, suggesting that cardiac function was impaired in the ICP fetus^[10]^. With the increased Tei index, the incidence of adverse perinatal outcomes, such as preterm birth, cesarean section, detection of abnormal fetal hearts, meconium contamination in amniotic fluid, low birth weight infants, and neonatal transfer to intensive care units, was increased^[11,12]^.

Previous studies mainly analyzed the relationship between maternal bile acid levels and fetal heart injury using fetal umbilical cord blood levels of myocardial enzymes, troponin levels, fetal electrocardiograms, and fetal heart color Doppler ultrasound. However, the results were varied and conclusions were not uniform, and few studies focused on the outcome of fetal heart injury after birth and its impact on the newborns. It has been reported that ICP patients and their babies may have an increased risk of developing heart disease^[15,16]^. Our study has focused on cardiac injury in the neonatal period. We analyzed the relationships between myocardial enzymes, ECG, and ultrasound echocardiography (UCG) of one-day-old neonates with maternal bile acid levels to study cardiotoxicity of ICP. The influence of the bile acid levels on the prognosis of neonatal cardiac trauma was analyzed by comparing myocardial enzymes from the above newborns five days after birth.

This study further confirmed that bile acid toxicity and the occurrence of adverse events in ICP were mainly related to bile acid damage to the fetal heart. In the treatment of ICP, emphasis should be placed on reducing the myocardial toxicity of bile acid to the fetus to avoid the occurrence of adverse events. The results of present study may provide theoretical support for this.

## Materials and Methods

### Patients and methods

#### Screening flowchart

The cases in our study included patients who gave birth in our hospital from September 10 2018 to June 30 2021, with their neonates transferred to the department of Pediatrics after birth immediately, excluding patients with multifetal pregnancies, diabetes, hypertension disorders, maternal/fetal heart disorders, and other liver disorders (Fig. 1). Patients’ characteristics of the three groups are listed in Table 1. The hospital numbers could identify individual participants during or after data collection. All data in ptesent study were fully anonymized before we accessed them and the Ethics Committee of Zhengzhou University had waived the requirement for informed consent.

**Table 1.**
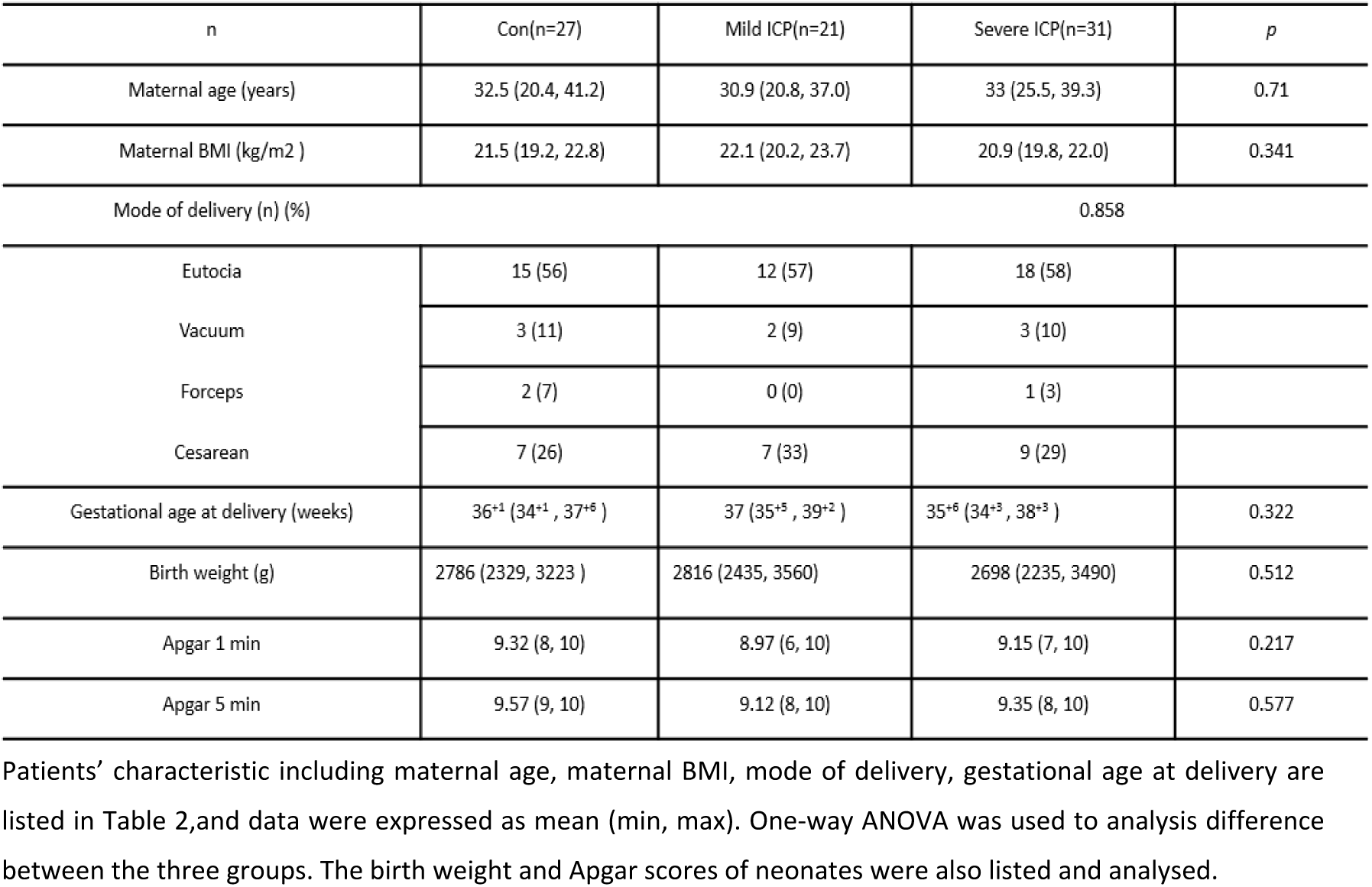
Patients’ characteristics of the three groups.

**Fig. 1.**
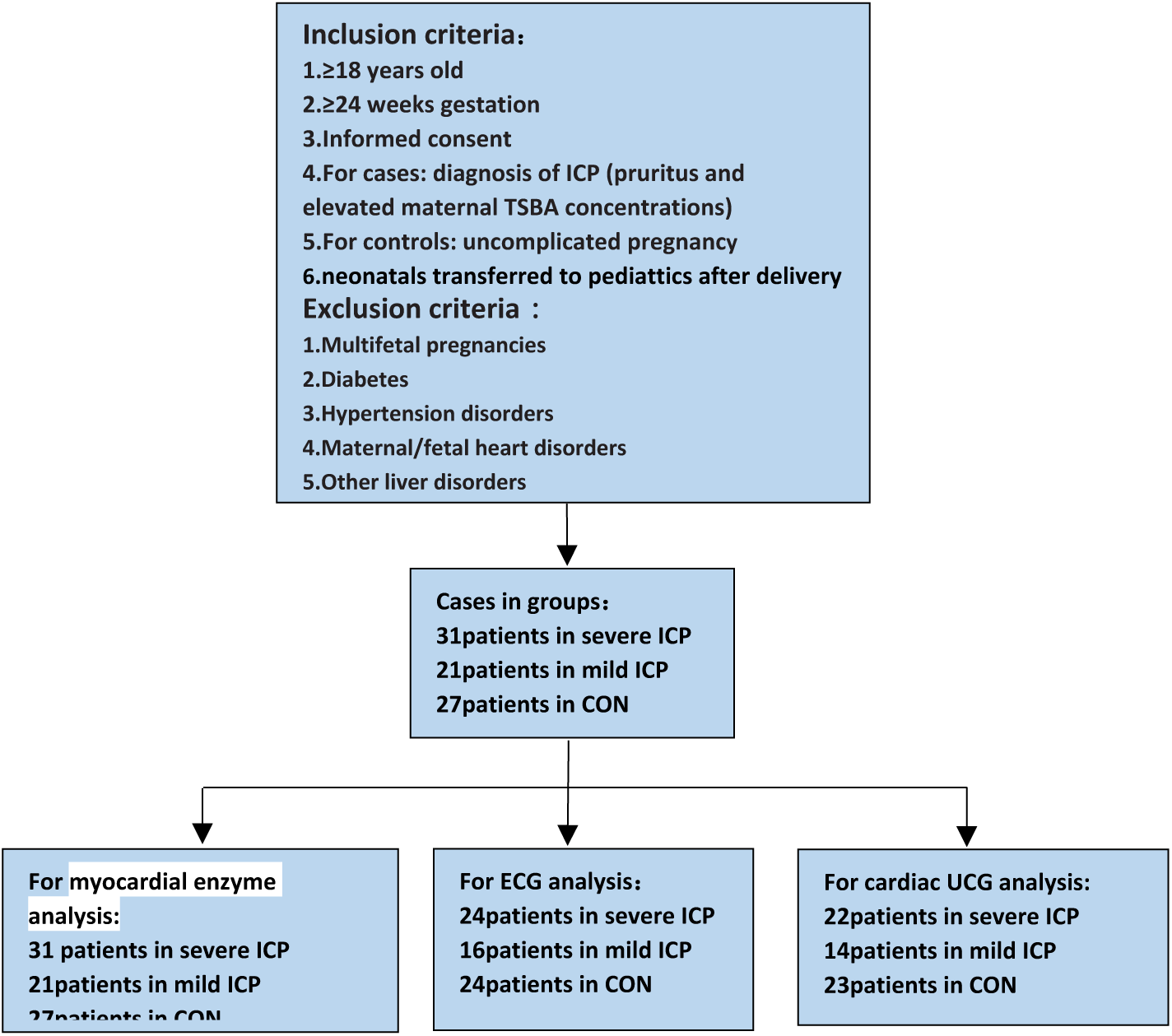
Flowchart depicting numbers of patients in each group analyzed. CON, control; ECG, electrocardiogram; UCG, ultrasound echocardiography; ICP, intrahepatic cholestasis of pregnancy; TSBA, total serum bile acids.

### Analysis methods

Bile acid levels and liver enzyme levels of pregnant women were recorded. According to the above exclusion criteria, 25 patients with mild ICP,31 patients with severe ICP and 27 normal control patients were selected and divided into mild ICP, severe ICP and normal control group. Neonatal cardiac parameters including myocardial enzymes[creatine kinase(CK), creatine kinase isoenzyme MB (CK-MB), lactate dehydrogenase(LDH), α-hydroxybutyric acid(HBDB)], ECG(heart rate, PR interval, QRS interval, QTc interval) and UCG[left ventricular ejection fraction (EF), left ventricular fractional shortening (FS)] were measured on the first day of birth and the fifth day of birth, respectively. Data analysis was performed using GraphPad Prism8.0 software(La Jolla, CA). One-way ANOVA was to analyze the difference of myocardial enzymes, ECG and UCG in the three groups, and a t-test was to compare the difference between each group. *P* < 0.05 was considered a significant difference. STATA16 (StataCorp, College Station, TX) was used to determine the correlation between the meaningful parameters and bile acid levels, and the influence of gestational days and delivery mode on the results. *P*<0.05 was considered a significant correlation.

## Results

### CK and CK-MB of one-day-old neonates were significantly elevated in ICP group

Peak concentrations of maternal total serum bile acid(TSBA) during pregnancy were used to classify women as mild (10–39 µmol/L) ICP1severe (≥40 µmol /L) ICP and CON group(<10µmol /L). Circulating levels of the liver-derived enzyme alanine aminotransferase were significantly higher in ICP group than thouse in the CON group, and those in the severe ICP group were significantly higher than those in the mild ICP group (Fig 2B), consistent with the differences in levels of TSBA. The CK and CK-MB levels were significantly higher in the severe ICP group compared with the CON group, but there was no significant difference between the mild ICP and CON groups (Fig. 2C, 2D). The levels of LDH, HBDB were similar among the three groups. The levels of myocardial injury indicator CK and myocardial injury-specific indicator CK-MB were significantly different between the groups.

**Fig. 2.**
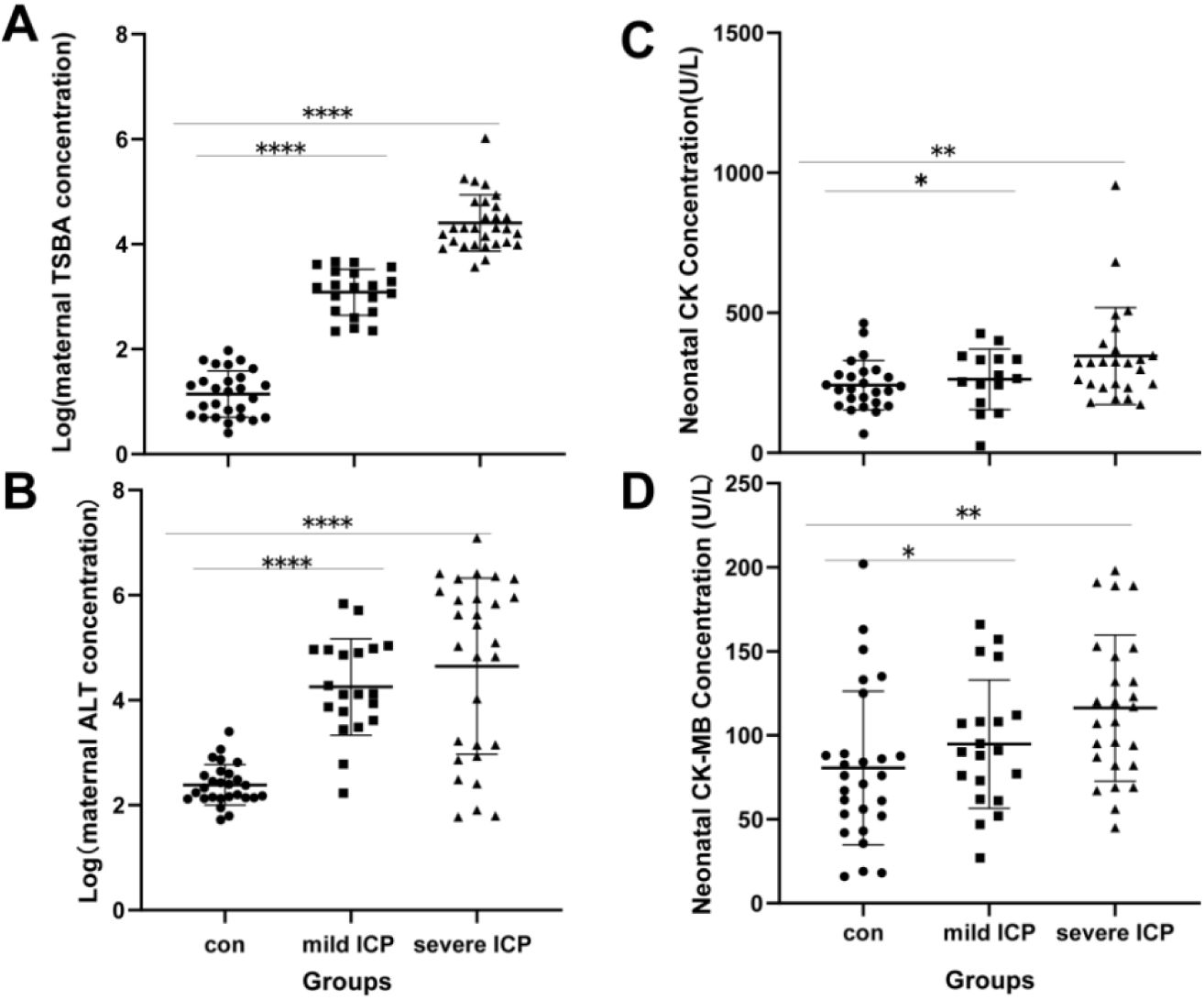
A, shows TSBA levels of pregnant women, with significant differences (ANOVA) among the three groups: con, control group; mild intrahepatic cholestasis of pregnancy, mild ICP group;severe intrahepatic cholestasis of pregnancy, severe ICP group. B, shows the serum levels of liver enzyme alanine aminotransferase (ALT) in pregnant women, with significant differences among the three groups, consistent with the findings of bile acid levels. C, shows CK levels in one-day-old neonates. ANOVA showed a significant difference, F = 4.292, P = 0.018. There was a significant difference between the severe ICP and con groups, but no difference between the mild ICP and con groups. D, shows significant differences (ANOVA) in CK-MB levels in one-day-old newborns (F = 4.599, P = 0.0133). The severe ICP group had significantly higher CK-MB levels than the mild ICP group, but there was no difference between the mild ICP and con groups. ****P < 0.001, **P < 0.01, *P > 0.05.

### QRS interval of neonatal ECG was significantly longer in severe ICP group

The one-way ANOVA findings showed that the diffierence of the QRS interval was significant, but the PR, QTc, and heart rate were not meaningfully different between the groups. The QRS interval of the severe ICP group was significantly longer than that of the mild ICP and CON groups, while there was no significant difference between the mild ICP and CON groups (Fig. 3A). These results indicated that the incidence of abnormal cardiac function, such as ventricular block, was higher in neonates from mothers with severe ICP. There were no significant differences in PR intervals, QTc intervals, and heart rates among the three groups (Fig. 3B–D).

**Fig. 3.**
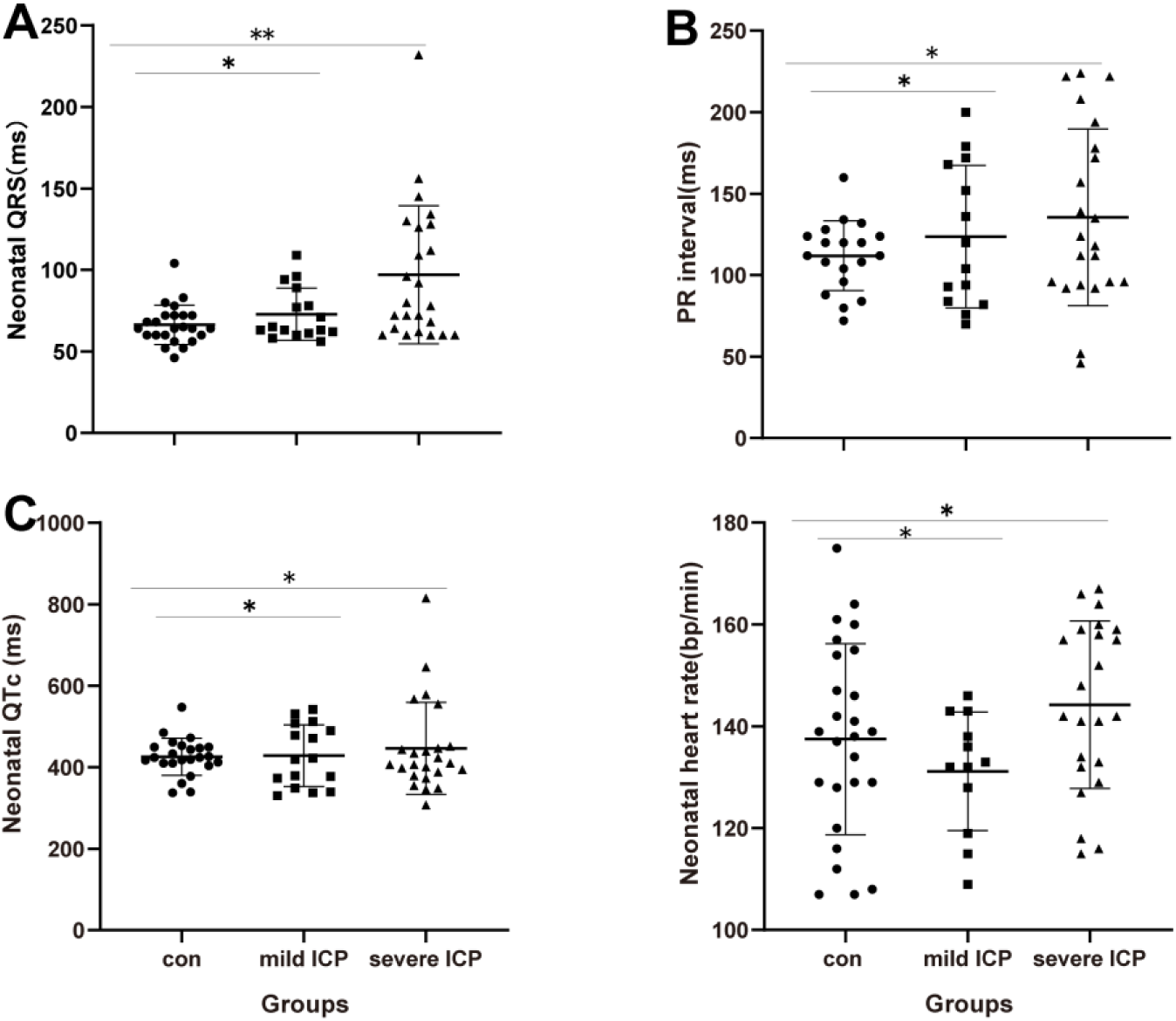
One-way ANOVA analysis of electrocardiograms (ECGs) of different neonatal groups. The ANOVA findings show differences in ECG recordings among the three groups, in particular significant differences in the QRS interval. A, The ECG QRS interval in one-day-old neonates showed a significant difference (ANOVA, F = 7.079, P = 0.001) between the three groups, with a larger interval in the severe ICP group compared with the mild ICP and con groups, noting that there was no significant difference between mild ICP and con groups. B–D, show ANOVA findings for the PR intervals, QTc intervals, and heart rates of one-day-old newborns in the three groups, respectively, and there were no meaningful differences. **P < 0.01, *P > 0.05.

### EF and FS of neonates from mothers with severe ICP were significantly lower than others

One-way ANOVA results indicated significant differences between the EF and FS measurements among the three groups. The EF was significantly lower in the severe ICP group than in the CON group, but there was no significant difference between the mild ICP and CON groups (Fig. 4A). The FS was significantly lower in the severe ICP group compared with the mild ICP and CON groups, while there was also no significant difference between the mild ICP and CON groups (Fig. 4B). The EF and FS were significantly different, indicating that ICP was related to the left ventricular ejection fraction and myocardial contractility of one-day-old neonates, and high concentrations of bile acid may reduce the left ventricular ejection fraction and myocardial contractility of one-day-old neonates.

**Fig. 4.**
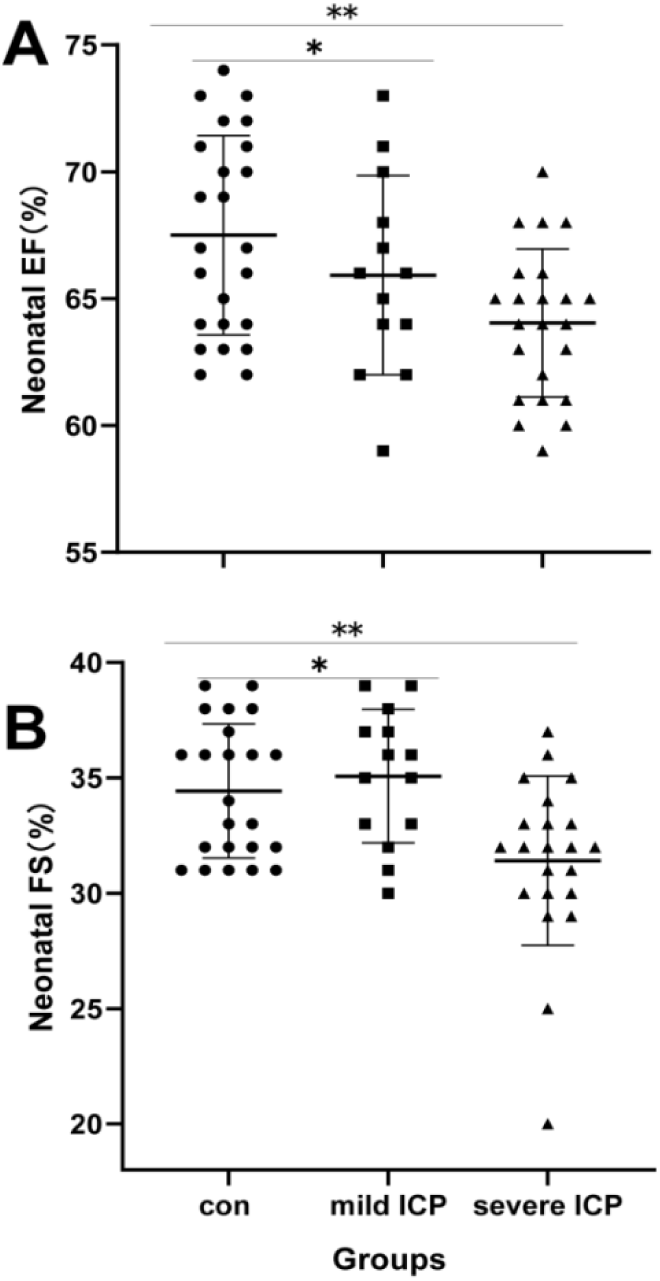
ANOVA analysis showed significant differences in color Doppler cardiac ultrasound indexes between the different groups. A, ANOVA of 1-day-old neonatal left ventricular ejection fraction (EF) showed a significant difference between the groups (F = 5.515, P = 0.0065). The EF was significantly lower in the severe intrahepatic cholestasis of pregnancy (ICP) group than the control (con) group (P = 0.0014). There were no significant differences between the severe ICP and mild ICP and between mild ICP and con groups. B, Analysis showed significant differences in left ventricular fractional shortening (FS) among the three groups (F = 7.304, P = 0.0015). The FS was significantly lower in the severe ICP group compared with the mild ICP (P = 0.0034) and control groups (P = 0.0037), while there was no significant difference between the mild ICP and control groups. **P < 0.01, *P > 0.05.

### There was a meaningful correlation between neonatal CK-MB, FS and maternal TSBA levels

The pwcorr and sig commands of STATA16 were used to analyze the correlation between neonatal cardiac parameters and maternal TSBA levels. The results showed a linear correlation between serum ALT and TSBA levels, and between CK-MB and TSBA levels in all three groups (Fig. 5A–C). There was no significant correlation between FS and TBSA levels. Further analysis showed that FS did not correlate with TSBA levels in the CON group (Fig. 5D), but was negatively correlated withTSBA in the mild and severe ICP groups (Fig. 5E). These results showed that myocardial contractility was significantly decreased in the severe ICP group, consistent with the differences in FS between the severe ICP and the other two groups. However, QRS, EF, and CK showed no obvious linear correlation with bile acid levels. These results indicated that CK-MB and FS of one-day-old neonates were affected significantly by maternal elevated TSBA. We propose that a maternal high concentration of TSBA caused neonatal myocardial damage and decreased myocardial contractility.

**Fig. 5.**
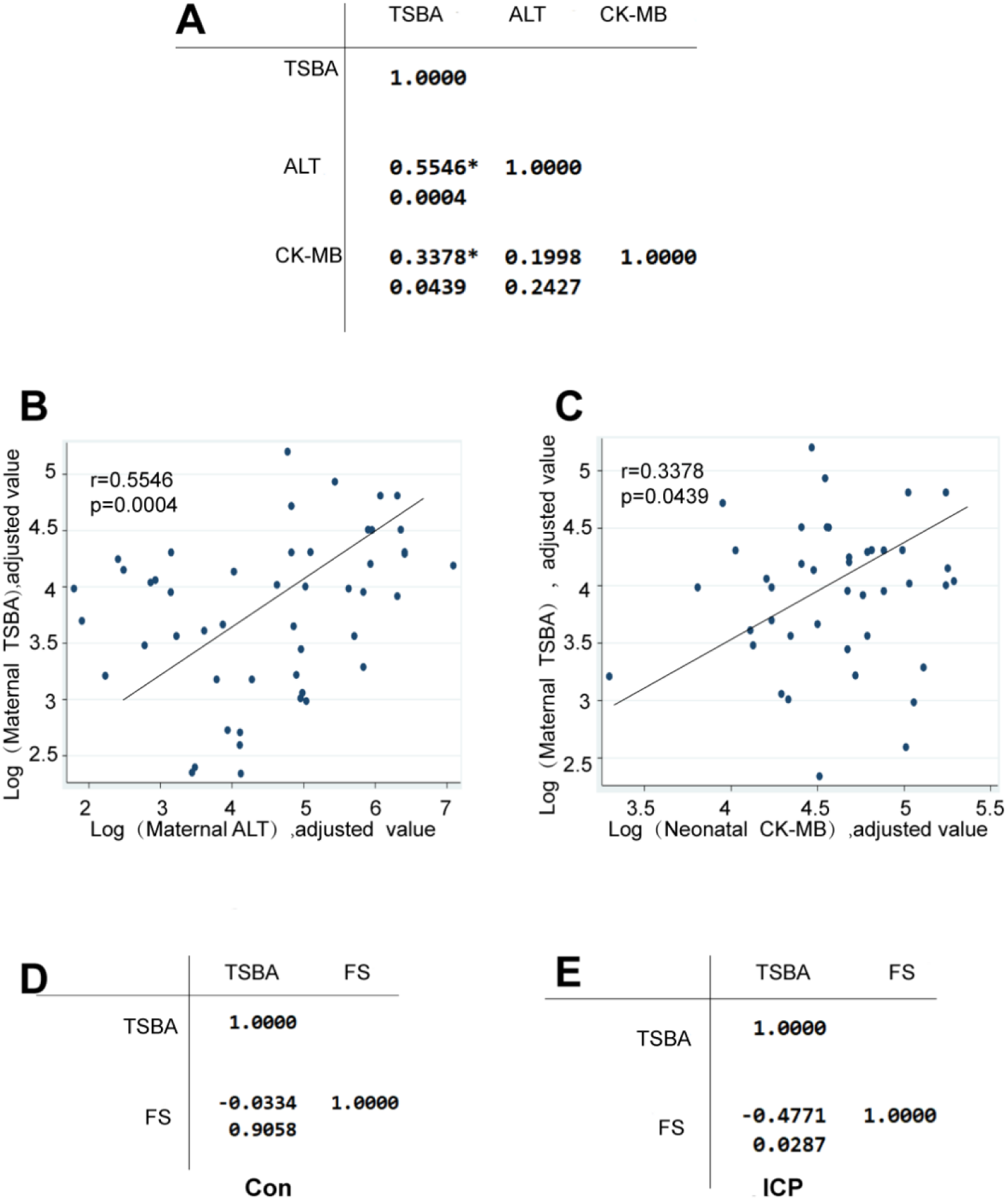
Correlation analysis of creatine kinase isoenzyme-MB (CK-MB), alanine aminotransferase (ALT), and left ventricular fractional shortening (FS) with serum bile acid levels. The pwcorr and sig commands of STATA16 were used to analyze the correlation between ALT, CK, CK-MB, QRS interval, EF, FS, and TSBA levels. A, shows a significant linear correlation between ALT and TSBA (r = 0.5546, P = 0.0004), and between CK-MB and TSBA levels (r = 0.3378, P = 0.0439). B and C, show scatter plots of the correlation between ALT and total TSBA levels, and between CK-MB and TSBA levels, respectively. D, shows there was no correlation between FS and TSBA in the control groups (r = −0.0334, P = 0.9058). E, shows a significant negative correlation between FS and TBSA in the ICP groups (r = −0.4771, P = 0.0287).

### The relationship of neonatal cardiac injury and maternal TSBA was unrelated to the days of gestation

Days of gestation has a decisive effect on neonatal outcomes and are correlated with neonatal phylogeny. To analyze the effect of gestational days on the relationship between bile acid levels and cardiac function, we used STATA16 to analyze the relationship between days of gestation and indicators of cardiac injury. The results showed that there was no significant correlation between CK-MB and days of gestation and total serum bile acid levels in the control group(Fig. 6A,B). In the ICP group, there was a significant positive correlation between CK-MB and total bile acid levels, and a significant negative correlation between CK-MB and days of gestation(Fig. 6C,D). Therefore, after excluding ICP, there was no correlation between days of gestation and CK-MB. In the ICP group, days of gestational were significantly negatively correlated with CK-MB levels, and total bile acid levels were positively correlated with CK-MB. We propose that a high concentration of bile acid plays a key role in myocardial injury and is directly related to myocardial injury indicators. Further analysis of ECG and UCG indicators of cardiac function also found that the FS and bile acid levels had no obvious correlation with days of pregnancy in the control group, but, in the ICP group, FS was positively correlated with days of pregnancy and exhibited a significantly negative correlation with bile acid levels. That is, ruling out the ICP factor, FS had no correlation with days of pregnancy. However, in the ICP group, the higher the bile acid levels and the shorter the period of gestation, the lower the measured FS, which represents myocardial contractility. There was no significant correlation between QRS with days of pregnancy both in the control and ICP groups.

**Fig. 6.**
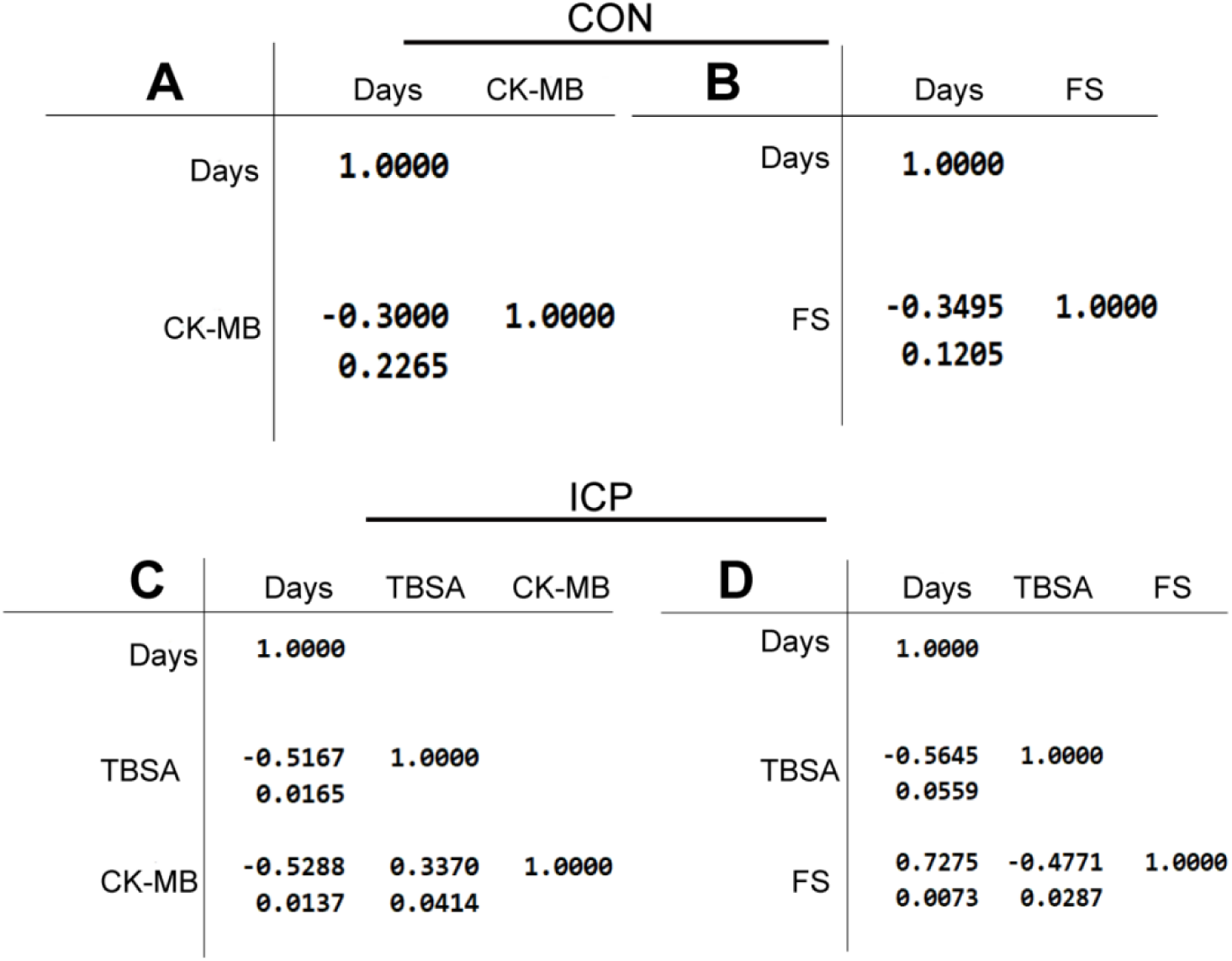
Correlation analysis of days of gestation with creatine kinase isoenzyme-MB (CK-MB) or left ventricular ejection fraction (FS). A and C, show that CK-MB, FS, and days of gestation had no significant correlations in the control group. B, shows that in the intrahepatic cholestasis of pregnancy (ICP) group, CK-MB was negatively correlated with days of gestation (r = −0.5288, P = 0.0137), and CK-MB was positively correlated with bile acid levels (r = 0.337, P = 0.0414). D, shows FS was positively correlated with days of gestation (r = 0.7074, P = 0.0068) and negatively correlated with bile acid levels (r = −0.4771, P = 0.0287) in the ICP group.

### The relationship of neonatal cardiac injury and maternal TSBA was unrelated to the delivery mode

The delivery mode may have some influence on newborn outcomes. To analyze the effect of delivery mode on the relationship between bile acids and heart damage, we divided all patients into a cesarean section group and a vaginal delivery group according to delivery mode. Analysis using the t-test showed no significant differences of CK-MB between the cesarean section and vaginal delivery groups(Fig. 7A). Similarly, FS had no significant differernces between the two groups. That is, delivery mode did not affect the analysis of CK-MB and FS in the current study(Fig. 7B).

**Fig. 7.**
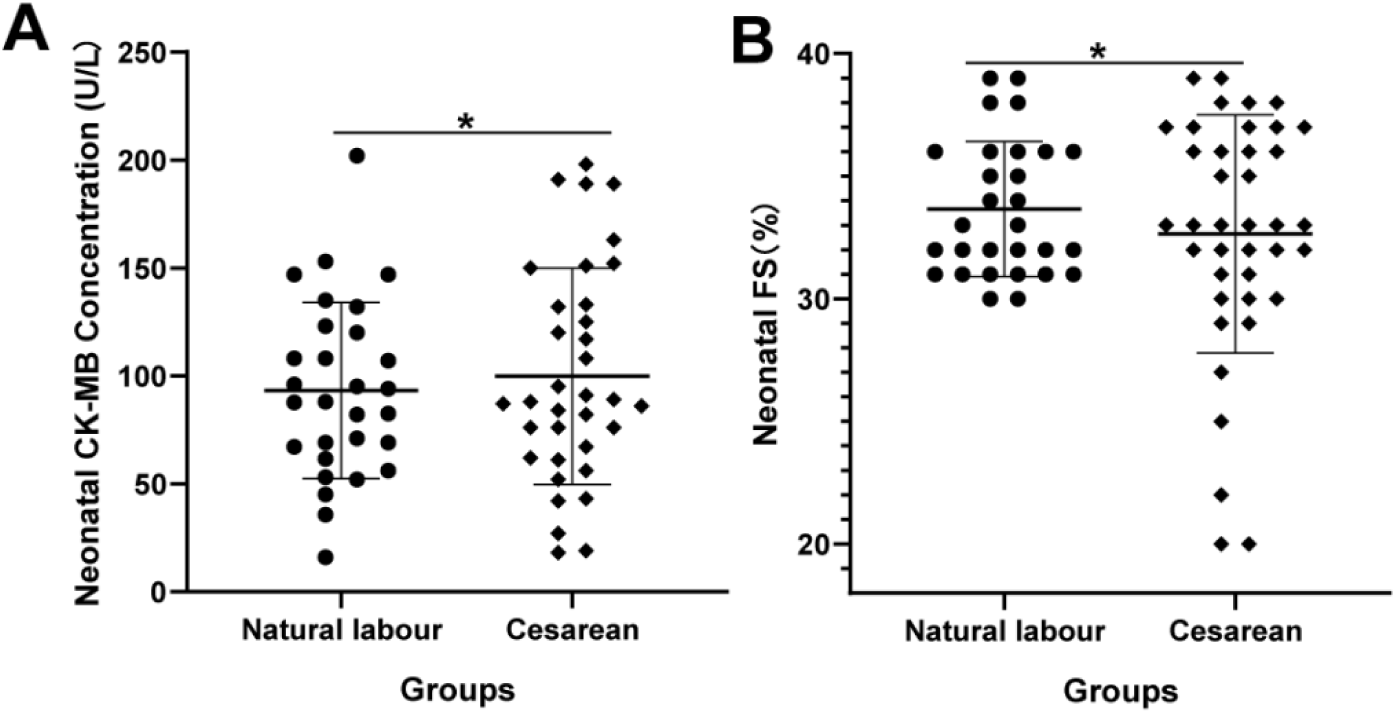
Analysis of variance of creatine kinase isoenzyme-MB (CK-MB) and left ventricular ejection fraction (FS) measurements for neonates from cesarean section and vaginal delivery groups. A, shows that there was no significant difference in CK-MB levels between the cesarean section and vaginal delivery groups. B, shows no significant difference in FS between the cesarean section and vaginal delivery groups. *represents P > 0.05.

### There was no difference in myocardial markers of five-day-old neonates between the ICP and CON groups

To determine whether the influence of TSBA on neonatal injury was persistent, we compared CK and CK-MB levels between the ICP and CON groups at the fifth day after birth. The t-test showed no significant differences between the two groups. Five days after birth, there were no differences in myocardial indicators between the ICP and CON groups(Fig. 8A,B). The myocardial indicators of the ICP group may quickly decrease to normal levels after the newborn being released from high concentrations of TSBA, resulting in no differences between the ICP and CON groups.

**Fig. 8.**
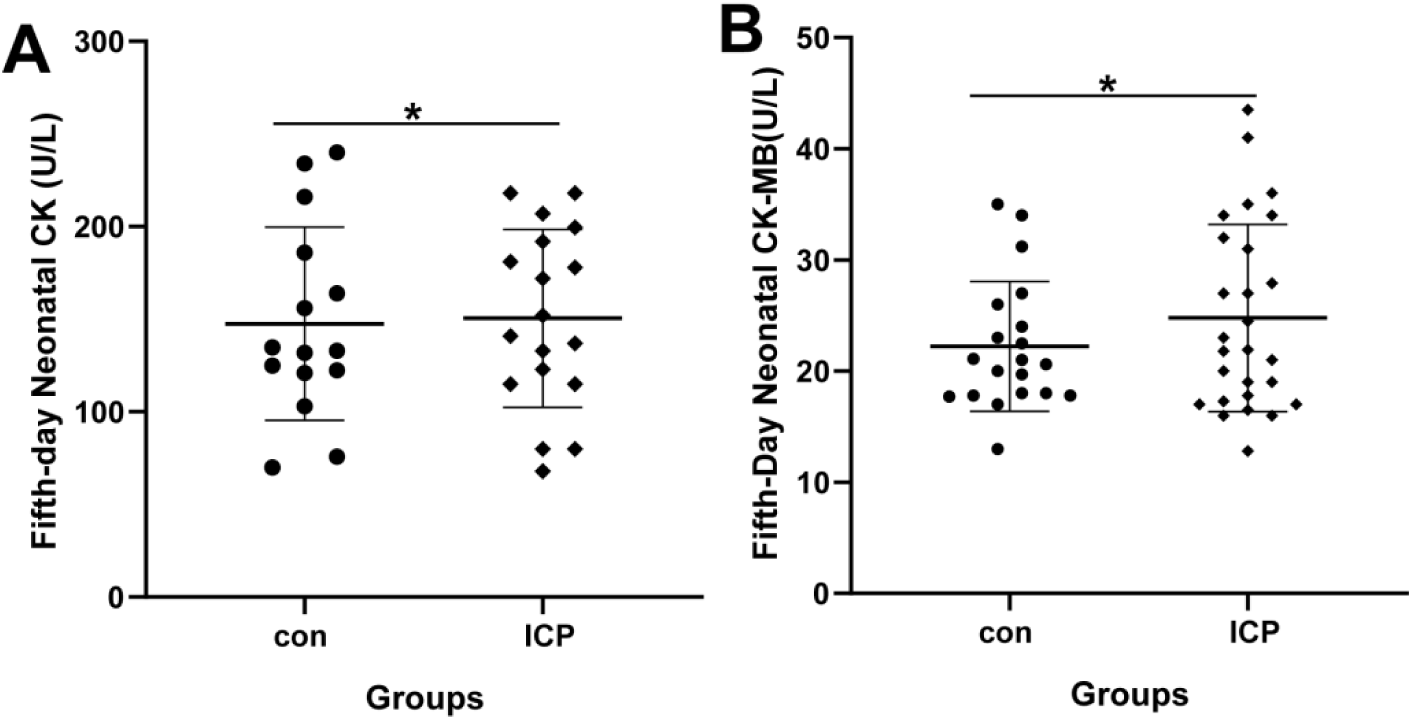
One-way ANOVA analysis of creatine kinase (CK) and CK isoenzyme-MB (CK-MB) levels of five-day-old infants from the control (con) group and intrahepatic cholestasis of pregnancy (ICP) group. A, shows no significant difference in CK levels between the con and ICP groups. B, shows no significant difference in CK-MB levels between the con and ICP groups. *P > 0.05.

## Discussion

At present, it is believed that the causes of sudden fetal death in patients with ICP mainly include insufficient oxygen supply from the placenta caused by placental villus space stenosis, amniotic fluid pollution, and fetal heart injury^[1]^. Fetal heart injury has emerged as a new factor for ICP stillbirth in recent years. Actually, sudden fetal death may result from the combined actions of multiple factors, such as fetal myocardial damage, fetal arrhythmia causing by the myocardial damage, inadequate placental oxygen supply aggravating the myocardial damage, contractions further reducing placental delivery of oxygen, and amniotic fluid contamination leading to fetal hypoxia. In all these factors or several factors under the joint action may lead to sudden fetal death in utero^[13,14]^. Fetal myocardial injury may play a key role in this process.

In this study, Myocardial enzymes, ECG, and cardiac UCG indicators of one-day-old neonates were measured(Table2), and significant differences between CON, mild ICP, and severe ICP groups were determined using one-way ANOVA. The current work showed that CK, CK-MB, QRS interval, and FS measurements were significantly different between the severe ICP and control groups, as well as between the severe ICP and mild ICP groups. Neonatal heart rate, PR interval and QTc interval showed no significant differences between these groups, findings that were not completely consistent with other reported fetal studies. Further analysis showed that there was a clear correlation between levels of CK-MB, an specific indicator of myocardial injury, and bile acid levels in all patients and groups, while there was no significant correlation between the non-specific myocardial enzyme CK and bile acid levels. There was no significant correlation between FS and bile acids or the liver enzyme ALT in the control group, but there was a significant negative correlation between FS and bile acids in all ICP patients. The QRS intervals were not significantly associated with bile acid levels in all patients or groups. In summary, the neonatal cardiac injury indicators associated with ICP were mainly the increased levels of CK-MB and the reduced FS, suggesting that the effects of bile acids on the neonatal heart mainly involved damage to the myocardial cells and weakening of myocardial contractility. Days of gestation are directly related to neonatal outcomes. We further study these datas to analyze the effect of gestational days on the relationship between bile acid levels and cardiac function. The current analysis of the above-mentioned indicators (CK, CK-MB, FS, and QRS) found no significant correlations with days of gestation in the control group. However, in the ICP group, CK-MB levels were positively correlated with bile acids and negatively correlated with days of gestation, and FS was negatively correlated with bile acids and positively correlated with days of gestation. The lower the level of bile acids, the longer the period of gestation in ICP patients, leading to reduced neonatal heart damage. Additionly, the delivery mode may affect the Apgar score and cardiac function of newborn babies. Therefore, this study investigated that effect of cesarean section and vaginal delivery on CK-MB and FS measurements. Analysis of the ICP group showed that there were no significant differences in CK-MB and FS levels in the cesarean section and vaginal delivery groups.

**Table 2.** One-way ANOVA analysis of maternal TSBA, ALT and cardiac parameters of one-day-old neonates.

| groups | CON | Mild ICP | Severe ICP | *p* |
| --- | --- | --- | --- | --- |
| TSBA (mmol/L) | 3.46 ± 0.30 | 23.77 ± 2.12 | 96.40 ± 13.62 | <.001 |
| ALT (U/L) | 11.73 ± 1.022 | 100.76 ± 19.96 | 260.27 ± 51.71 | <.001 |
| CK (U/L) | 241.64 ± 17.59 | 262.74 ± 27.83 | 346.05 ± 34.56 | .0180 |
| CK-MB (U/L) | 80.50 ± 8.80 | 94.7 ± 8.53 | 116.2 ± 8.53 | .0133 |
| LDH (U/L) | 428.6 ± 24 | 476.6 ± 16.9 | 417.4 ± 14.10 | NS |
| HBDB (U/L) | 280.5 ± 18.83 | 340.4 ± 23.25 | 297.0 ± 13.01 | NS |
| QRS (ms) | 66.33 ± 2.473 | 72.81 ± 4.02 | 97.00 ± 8.63 | .001 |
| QTc (ms) | 425.7 ± 9.28 | 428.6 ± 18.87 | 446.2 ± 23.03 | NS |
| PR (ms) | 111.9 ± 4.91 | 123.6 ± 11.68 | 135.5 ± 11.57 | NS |
| FS (%) | 34.43 ± 0.60 | 35.07 ± 0.77 | 31.41 ± 0.78 | .0015 |
| EF (%) | 67.50 ± 0.80 | 65.92 ± 1.09 | 64.04 ± 0.61 | .0065 |

CON: control; ICP: intrahepatic cholestasis of pregnancy; TSBA: total serum bile acid; ALT: alanine aminotransferase; CK: creatine kinase; CK-MB: creatine kinase isoenzyme MB; LDH: lactate dehydrogenase; HBDB: a-hydroxybutyric acid; QRS and QTc: ECG parameters; PR: pulse rate; FS: left ventricular fractional shortening; EF: ejection fraction. Results are expressed as means ± S.E.M.

To analyze the outcome of neonatal heart injury in the ICP group, we reviewed the myocardial enzyme indicators at the 5th day of birth. One-way ANOVA findings showed that CK, CK-MB, LDH, and α-hydroxybutyrate dehydrogenase levels were similar in the three groups. To some extent, these findings indicated that the outcome of myocardial injury in one-day-old neonates from patients with ICP was consistent with that from normal patients. The fetal heart injury caused by bile acids may automatically recover in the time after the fetus was separated from the high bile acid environment of the mother. These findings also indicated that the myocardial injury of fetus and 1-day-old neonates were closely related to the high bile acid levels. However, because of the lack of ECG and cardiac ultrasound analysis in 5-day-old newborns, as well as the lack of long-term follow-up results, the present study cannot rule out long-term effects of bile acids on newborns, and whether they will exhibit an increased risk of heart disease.

The current results, combined with previous studies, show that a high concentration of bile acids cause fetal heart damage, which is mainly characterized by down-regulation of troponin I, prolongation of the PR interval, and atrioventricular transmission block at the fetal stage^[5,8]^. In 1-day-old neonates, the main features of cardiac damage were increased specific myocardial enzyme CK-MB, reduced FS, myocardial damage, premature ventricular beats, and bundle branch block. However, due to the small number of participants in present study and the lack of long-term follow-up studies, further analysis of larger and longer studies is needed to draw firm conclusions.

## Conclusions

In this study, the cardiotoxic effect of maternal TSBA was demonstrated in neonates, and the outcome of heart injury in neonates from patients with ICP was analyzed. In the future, basic experiments should explore the mechanism of myocardial injury by biel acid, as well as the key molecules and signaling pathways involved, to provide new avenues for the clinical treatment of ICP and the prevention of stillbirth in pregnant women with ICP.

## Fundings

Fund Source: Science and Technology Key Projects of Henan Province (Fund No.2018020189;Fund No. LHGJ20190363)

## Author contributions

All authors contributed significantly to this work. Junnai Wang: Optimized the analytical methods, extracted and analyzed the data, and wrote the manuscript; Wei Shi: collected the samples and conducted experiments; Weiwei Lun: Designed the research, analyzed the data and wrote the manuscript. All authors read and approved the final manuscript.

## Disclosure of interest

The authors declare that they have no competing interest.

## Acknowledgments

We thank the FUNDS (2018020189 and LHGJ20190363) from Science and Technology Key Projects of Henan Province for supporting this research. We also thank Liwen Bianji (Edanz) (www.liwenbianji.cn/), for editing the English text of a draft of this manuscript.

## Notes

### Competing Interest Statement

The authors have declared no competing interest.

